# Maximum likelihood method quantifies the overall contribution of gene-environment interaction to continuous traits: an application to complex traits in the UK Biobank

**DOI:** 10.1101/632380

**Authors:** Jonathan Sulc, Ninon Mounier, Felix Günther, Thomas Winkler, Andrew R. Wood, Timothy M. Frayling, Iris M. Heid, Matthew R. Robinson, Zoltán Kutalik

**Author notes:** These authors contributed equally to this work (Jonathan Sulc, Ninon Mounier).

## Abstract

As genome-wide association studies (GWAS) increased in size, numerous gene-environment interactions (GxE) have been discovered, many of which however explore only one environment at a time and may suffer from statistical artefacts leading to biased interaction estimates. Here we propose a maximum likelihood method to estimate the contribution of GxE to complex traits taking into account all interacting environmental variables at the same time, without the need to measure any. This is possible because GxE induces fluctuations in the conditional trait variance, the extent of which depends on the strength of GxE. The approach can be applied to continuous outcomes and for single SNPs or genetic risk scores (GRS). Extensive simulations demonstrated that our method yields unbiased interaction estimates and excellent confidence interval coverage. We also offer a strategy to distinguish specific GxE from general heteroscedasticity (scale effects). Applying our method to 32 complex traits in the UK Biobank reveals that for body mass index (BMI) the GRSxE explains an additional 1.9% variance on top of the 5.2% GRS contribution. However, this interaction is not specific to the GRS and holds for any variable similarly correlated with BMI. On the contrary, the GRSxE interaction effect for leg impedance 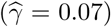 is significantly (*P* < 10^−56^) larger than it would be expected for a similarly correlated variable 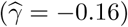. We showed that our method could robustly detect the global contribution of GxE to complex traits, which turned out to be substantial for certain obesity measures.

## 1 Introduction

Genome-wide association studies (GWAS) have been instrumental in the discovery of tens of thousands of genetic variants (mainly single nucleotide polymorphisms, SNPs) associated with complex traits and diseases^1^. While dozens or even hundreds of SNPs have been found to be associated with each trait, the effect contributed by each individual marker is typically very small^2^. In addition to their marginal effects, many genetic factors are suspected to alter susceptibility to the effects of environmental factors on the trait. The detection of these gene-environment (GxE) interactions has become possible with the large sample sizes of current GWAS and many methods have been proposed to achieve this. However, most of these methods assume that the genetic marker(s) and the interacting environmental factor have been measured without noise, and often require that the outcome and/or environment be binary. While model misspecification may have a more limited impact in single SNP association analysis, this can result in biased estimation of the interaction term in GxE analysis^3^. For example, testing a noisy or dichotomised version of a continuous environmental variable can bias the estimation in an arbitrary direction depending on the dichotomisation threshold^4^. Observing the outcome variable on a transformed scale (i.e. where the effect of the genetic marker is not linear) is also liable to introduce bias in any direction and magnitude.

More advanced methods adapt a two-step procedure including a filtering first step based on either the marginal SNP-trait association^5,6^, SNP-environment association^7^, a mixture of the two (cocktail method)^8^ or their combination^9^. Mixed effect models have been proposed to evaluate the differences in heritability in subgroups based on environmental exposure^10^. All these methods, however, require that the environment be measured (accurately), whereas in most studies only some of the relevant environmental variables are reported. Furthermore, many potentially crucial factors may be difficult to define precisely or measure accurately, such as physical activity, accessibility of fast food, sleep, and diet which are all key factors in obesity and are suspected to interact with genetic risk. Even in cases where a potentially relevant factor has been measured accurately, it is impossible to determine whether any detected interaction is due to the tested variable or one of its correlates. For example, many environmental factors have been shown to interact with the genetic risk score for body mass index (BMI)^11^, such as physical activity^12^, alcohol consumption^13^, socio-economic status (as measured by the Townsend deprivation index)^14^, sugary drink consumption^15^, certain types of diet^16^, etc. Many of these environmental variables are correlated with one another and little is known about how these interactions relate to each other^13^.

Another approach which has been proposed is to detect GxE interaction based on differences in variance across genotype groups^17–19^. This has the advantage of accounting for all interacting environmental variables, without requiring their observation. However these methods were not designed to assess the extent of the interaction strength and are mostly restricted to single SNP analysis. In addition, these studies do not seek to account for general scale effects that are not specific to the genetic markers. Due to their low statistical power, variance tests have rather been used to improve power of classical GxE tests where the environment is measured and testable^20^. Others also proposed variance component analysis^21^, assuming that the environment is emerging as cumulative effect of multiple observed molecular phenotypes, which is less feasible for complex human traits and the method is not scalable to hundreds of thousands of samples.

In this work, we propose a method to establish statistical interaction between a genetic risk score (GRS) for a continuous outcome and all environmental variables. Furthermore, this method quantifies the total contribution of GxE to the trait variance beyond that of the GRS alone. The structure of the paper is as follows: First, we set up the normal linear interaction model and derive how to quantify the total GxE contribution. Next, we demonstrate through extensive simulations that any violations of the model assumptions (such as normality of the underlying environment and noise) do not introduce noticeable bias in the interaction parameter estimation. Further, we show that our bootstrap procedure produces close to nominal coverage probability of the produced confidence intervals regardless of the noise distribution. In addition, we propose an approach to determine whether the observed variance inflation is specific to the tested GRS or simply due to general heteroscedasticity. Finally, we apply our method to the GRSs of 32 continuous complex traits from the UK Biobank to reveal the contribution of GxE to their variability. The code implementing the algorithm is available in R and Matlab (https://github.com/zkutalik/GRSxE_software).

## 2 Methods

In the past, methods have been proposed to detect GxE based on variance heterogeneity of the outcome in different genotype groups^17^. These methods, like ours, do not need to observe the interacting environment as they treat the environment as a nuisance variable and integrate it out. This results in loss of statistical power as we only look at the consequence of such interaction in terms of the change in the outcome distribution as a function of the GRS value. The presence of GxE would lead to a different distribution of the outcome (principally characterised by increased variance) in higher genetic risk groups, which is captured by the fitted likelihood function. Our maximum likelihood approach does not require any arbitrary grouping of the population into subgroups according to their genetic predisposition and the interaction effect size is directly estimated.

### 2.1 Derivation of the likelihood function

Let us define a set of *n* individual and ***y*** ∈ ℛ^*n*^ denotes the observed outcome variable, ***e*** ∈ ℛ^*n*^ is an (unobserved) environmental factor and the ***g*** ∈ ℛ^*n*^ is the available GRS (or as a special case the genotype of a single SNP) in this sample. The central focus of our paper is to detect the presence of environmental influence that modifies the effect of the genetic risk score (***g***) on the outcome (gene-environment interaction) and to quantify the extent of this modification. Here, we are particularly interested in an abstract environmental variable (which captures the impact of several measurable exposures such as socio-economic status, physical activity, alcohol consumption, etc.) that modifies the genetic predisposition of a complex trait (***y***). The corresponding capital letters represent the random variables. We assume that the true underlying model is as follows

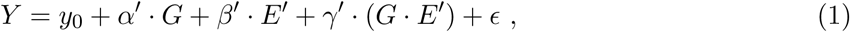

where, for notational simplicity, we assume that variables *Y, G* and *E*′ have zero mean and unit variance. The term *y*_0_ refers to the constant intercept. We allow for *E*′ and *G* being correlated (with correlation *δ*′) and for simplicity we assume linear relationship 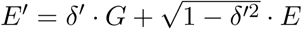, where *G* and *E* are independent and *E* has zero mean and unit variance. Hence *E* can be viewed and the part of the interacting environment *E*′ that is orthogonal to the genetic risk (*G*). The model can then be rewritten as

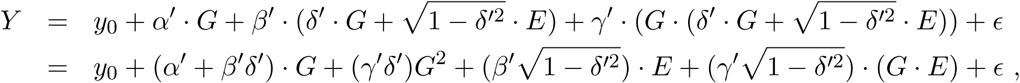

where *ϵ* is assumed to be normally distributed with zero mean. Note that due to the properties of *G, E* and *Y* all terms on the right hand side have zero mean, except *E*[(*γ*′ *δ*′)*G*^2^] = (*γ*′*δ*′), hence *y*_0_ = −(*γ*′*δ*′). Therefore, the model simplifies to

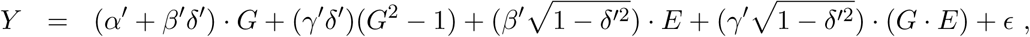

This model is equivalent (and can be reparameterized) to

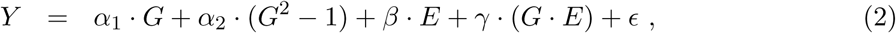

with 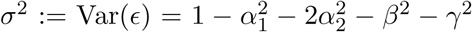. Here we defined *α*_1_ := *α*′+ *β*′*δ*′ as the observed linear genetic effect, *α*_2_ := *γ*′*δ*′ is the quadratic effect due to a non-zero interaction and *G - E*′ correlation, 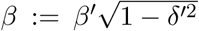 is the *G*-independent environmental effect and 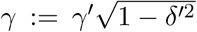 stands for the interaction effect between *G* and the *G*-independent environment. Note that we cannot distinguish between an interaction model with pure linear *G* term with correlated G-E (Eq. (1)) and a model with uncorrelated G-E, but with quadratic relationship between *Y* and *G* (Eq. (2)). Since the two models are mathematically equivalent, we will continue working with the latter parameterisation. Note that the model is parameterised such that the variance explained by the GxE term is simply *γ*^2^. If one wishes to recover the explained variance of the original *E*′ (i.e. *γ*′), it is given by 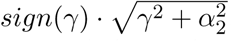.

We can write the density function of (*Y* |*G* = *g*) as

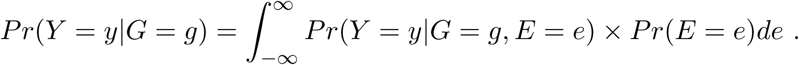

We assume that *E* and *ϵ* to be normally distributed, i.e. *Pr*(*E* = *e*) = *ϕ*(*e*) and *Pr*(*ϵ* = *e*) = *ϕ*(*e*), where *ϕ*(·) is the probability density function of the standard normal distribution. Therefore, the integral simplifies to

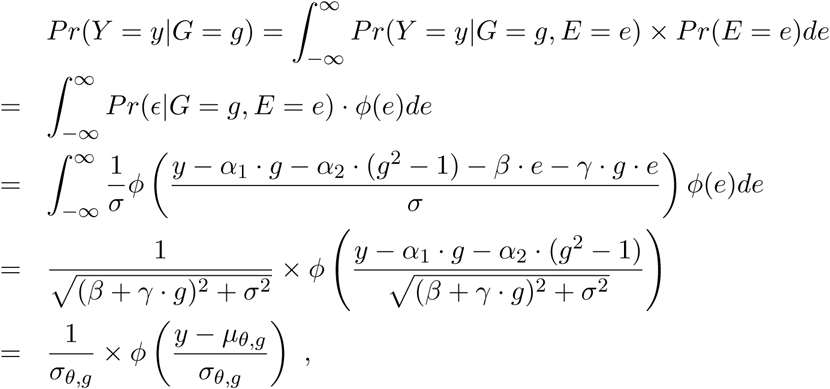

where *θ* = {*α*_1_, *α*_2_, *β, γ*}, *µ*_*θ,g*_ = *α*_1_ · *g* + *α*_2_ · (*g*^2^ − 1) and 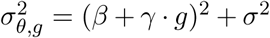. The likelihood function (for observed data (*Y, G*)) can be written as

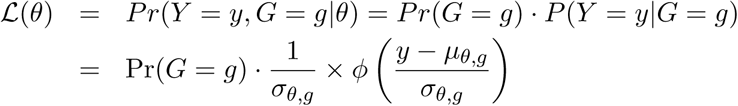

To estimate the underlying parameters, we need to maximise the log-likelihood function, i.e.

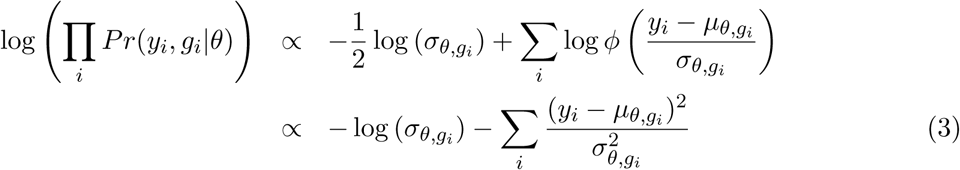

Note that the minimisation was constrained such that 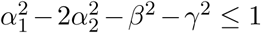 so that Var(*ϵ*) ≥ 0. Covariates, (e.g. age, sex, ancestry principal components) can be incorporated into the model by modifying *µ*_*θ,g*_ to *α*_1_ · *g* + *α*_2_ · (*g*^2^ − 1) + `C` · **c**, where `C` is the matrix with all the covariates listed as columns and **c** is the coefficient vector that will be estimated in the ML procedure.

### 2.2 Variance of parameter estimates

When the error term substantially deviates from normal distribution (the deviation limit depending on sample size), the variance of the parameter estimates (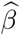 and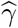) cannot be derived reliably from the log-likelihood function (Eq. (3)), e.g. by computing the Fisher information matrix or via likelihood ratio test. Instead, we performed 100 bootstrap sampling to obtain robust variance estimates. Note that this is preferable to a permutation procedure since – due to the special nature of our likelihood function – the estimator variance is larger under the null than under the true interaction model, hence it would lead to decreased power and conservative confidence interval. For real data application, when more than 10% of the bootstrapped data yielded estimates with Var(*ϵ*) = 0 (i.e. maximisation stuck at the boundary) the analysed trait was discarded to be on the safe side. To reduce computational time when testing large number of single SNPs (instead of one GRS), one can compute the Fisher information matrix to obtain standard error for the interaction estimates and use bootstrapping only for SNPs showing the most evidence for interaction. Note that when *γ* = 0, parameter *β* cancels out in the formula and hence not identifiable. For this reason, likelihood ratio test is not ideal to derive confidence interval for 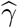.

### 2.3 Accounting for transformation of the outcome variable

It is possible that we do not directly measure (or model) the outcome, but only a transformed version of it, i.e. our observed data is {*y* = *f* (*z*), *g*} with the model *Z* = *α*_1_*G* + *α*_2_*G*^2^ + *βE* + *γ*(*G* × *E*) + *ϵ*. Such a transformation may induce general heteroscedasticity, which translates to mean-variance relationship that is not specific to any predictor of *Y*, i.e. any *X* with a non-zero correlation *r* with *Y* the *V ar*(*Y |X* = *x*) = *g*(*r, x*). However, we can test the specificity of the interaction effect identified through our variance modelling (Eq. 3) by simulating a counterfeit *G* (termed 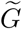) with properties similar to those of *G*. Specifically, we ensure it is similarly distributed and identically correlated to *Y* in terms of first and second moments 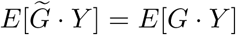 and 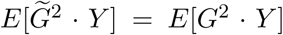, respectively). If the interaction obtained from applying the model described in Eq. 2 is not due to a transformation of the outcome, applying the same model to 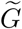 should yield no interaction. A similar interaction obtained using 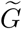 would indicate that the GxE we detect is not specific to *G* and is most likely due to observing the true underlying trait on a transformed scale.

To do so, we use the data (***g, y***) to estimate *E*[*G* · *Y*] = ***g***′ · ***y****/n* =: *b*_1_ and *E*[*G*^2^ · *Y*] = ***g***′^2^ · ***y****/n* =: *b*_2_. We choose to create a 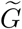 of the following form

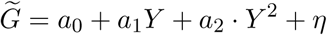

with *η* ∼ 𝒩 (0, *τ*^2^) and *E*[*η* · *Y*] = *E*[*η* · *Y* ^2^] = 0. Let us define *µ*_*i*_ := *E*[*Y*^*i*^] for *i* = 1, 2, …, 5 with *µ*_1_ = 0 and *µ*_2_ = 1. These can be estimated from the data as 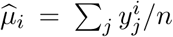. Note that if *Y* were normally distributed, *µ*_3_ = *µ*_5_ = 0 and *µ*_4_ = 3. To find *a*_0_, *a*_1_, *a*_2_ that satisfies 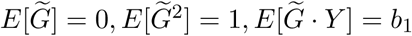 and 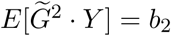 we need to solve the following equations

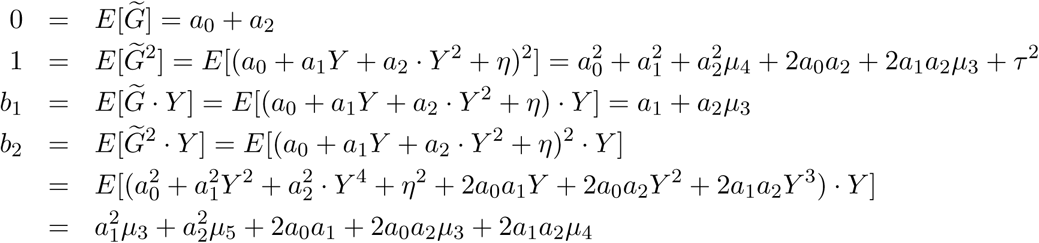

From the first equation we have *a*_0_ = −*a*_2_, the second yields 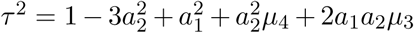 and the third equation gives *a*_1_ = *b*_1_ − *a*_2_*µ*_3_. Therefore, knowing *a*_2_ directly yields *a*_0_, *a*_1_ and *τ*.

Plugging these into the last equation gives

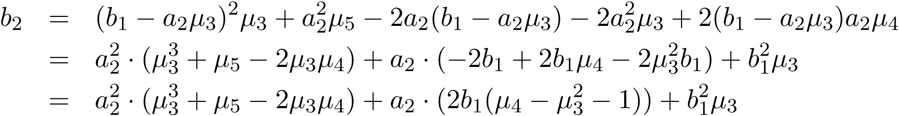

which is a second order polynomial in *a*_2_ and hence its solution is

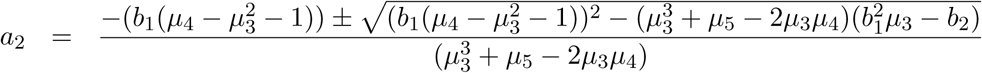

Having estimated *µ*_*i*_s and *b*_*i*_s from the data, we can obtain *a*_*i*_s and *τ*^2^, which can be used to simulate many counterfeit 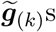. Finally, we fit the likelihood function (Eq. (3)) to this data 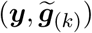 to generate a null distribution of 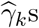. We then can compare the 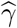 best fitting the true data (***y, g***) to the distribution of 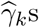 best fitting the counterfeit (genetic) data 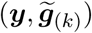 to test whether the observed interaction effect is specific to the GRS or common to any variable identically correlated with the outcome.

### 2.4 Simulation settings

We systematically explored the robustness of our method to various simulation settings. As a basic setting, we assume that the generated phenotype is governed by the interaction model defined in Eq. (2). To this end we simulated ***g*** ∼ 𝒩 (0, 1), ***e*** and *ϵ* The latter two were simulated from a Pearson distribution (R function `rpearson` / Matlab function `pearsrnd`), for which we set the first four moments (mean=0, variance =1, for skewness and kurtosis see below). In theory, violations of the normality assumption for *ϵ* and ***e*** could lead to biased estimation of the key parameters (*γ* and *β*). Therefore, we conducted extensive simulations including 21 different distributions with a wide range of values for skewness (*E*[*X*^3^] ∈ [0, 5]) and kurtosis (*E*[*X*^4^] ∈ [2, 27]) both for ***e*** and *ϵ* (Fig. S1). For these simulations we set other parameters to 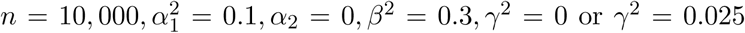 or *γ*^2^ = 0.025. We set slightly exaggerated effect sizes in order to keep the sample size relatively low, in order to save run time. For each parameter setting, we repeated the simulations 100 times. When these results are presented on boxplots, boxes mark the first (*q*_1_) second (*q*_2_) and third quartiles (*q*_2_) and the lower/upper whiskers are at *q*_1_ − 1.5 · (*q*_3_ − *q*_1_), *q*_3_ + 1.5 · (*q*_3_ − *q*_1_), respectively.

Next, we tested the impact of trait transformations (*f* (*t*) = *t*^*k*^, *k* = −1, 0, 1, 2, 3), where instead of observing/modelling (*Y, G*) directly, we observed (*f* (*Y*), *G*). Note that *k* = 0 represents the log-transformation. As the interaction effects are not the same on the transformed scale as on the original one, in these analyses our aim was only to distinguish between true and null interaction effects. Thus, we tested *γ* = 0 and *γ*^2^ = 0.05, while fixing other parameters at 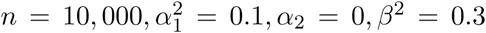. Furthermore, we tested the impact of combined violations of the model assumptions and simulated more data under all possible combination of transformations *f* (*t*) = *t*^*k*^(*k* = 0, 1, 2), skewness (*E*[*X*^3^] ∈ [0, 5]) and kurtosis (*E*[*X*^4^] ∈ [2, 27]) both for ***e*** and *ϵ*, resulting in further 126 (3 × 21 × 2) parameter combinations.

We then explored the power to discover GxE interactions for various combinations of realistic sample sizes (*n* = 10*K*, 20*K*, …, 100*K*) and interaction effects (*γ*^2^ = 0.2%, 0.4%, …, 2%). Since in a typical scenario one would test a few dozen outcome traits, we used a P-value threshold of 10^−3^ to establish power. For these simulations we set other parameters to 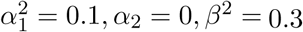. We have also compared power of our method working with unobserved *E* against linear interaction model with observed *E* under fixed setting of 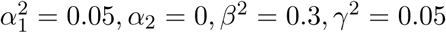 with all possible combinations of skewness and kurtosis for *E*.

In the simulations, the GRS (***g***) was assumed to have been measured without noise, while in real applications we can only estimate it. To examine whether using an estimate rather than the exact GRS results in estimation error, we simulated 1,000 independent genotypes for a sample of 10,000 participants. The phenotype was then generated based on a polygenic model, where the effects of SNPs are drawn from a normal distribution and explain in total 30% of the trait variance. We then added an environmental factor (*E*) and an interaction term (*GRS* × *E*) to this genetic component, explaining 30% and 10% of the trait variance, respectively. Finally, we added normally distributed noise, contributing the remaining 30% to the phenotypic variance. Note that we used here larger SNP and GxE effects, because as opposed to the other simulations, here we created GRS not only based on genome-wide significant SNPs, while all previous settings mimic real scenarios, where the GRS is composed of only genome-wide significant SNPs. Using the simulated phenotype and genotype data we derived GRS at different P-value thresholds. We computed two different GRSs, one based on ordinary least squares (OLS) and another one using the best linear unbiased predictor (BLUP). The emerging GRSs were then plugged into the likelihood function.

### 2.5 Application to complex continuous traits in the UK Biobank

To explore whether we can find evidence of interaction between GRS and environmental variables, we looked at 32 continuous complex traits, where the GRS—composed of all independent, genome-wide significant SNPs in the UK Biobank— explained at least 2% of the variance of that trait. For this we first selected SNPs with association P-value < 5×10^−8^ and pruned them based on distance, eliminating SNPs that lie within 1Mb vicinity of a stronger associated variant. We define an outcome trait continuous if the variable takes at least 100 different values in the UK Biobank sample. The analysis was restricted to a sub-sample of the UK Biobank comprising 378,836 unrelated, white British participants. We fitted the likelihood function both to the real data (*Y, G*) and 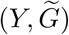 counterfeit data in order to assess whether the detected interactions are specific to the GRS or generally present due to scale issues. When measures were available from both left and right side of the body (e.g. left and right arm fat mass), we used only the right side measures due to extremely high correlation (> .95) between the left and right traits.

### 2.6 Single SNP analysis

Single SNP interaction testing differs in two aspects from the standard GRS analysis. On one hand, the expected interaction effects are much smaller and the bootstrapping procedure is not feasible for millions of SNPs genome-wide. To address the first point, we extended the simulations to smaller interaction (*γ*) values down to 0.02%) latter point. To reduce computational time when testing large number of single SNPs, one can use the likelihood ratio test to obtain standard error for the interaction estimates and use only bootstrapping for SNPs showing the most evidence for interaction.

## 3 Results

As outlined in Section 2, we calculate the overall contribution of GxE interaction between a fixed genetic factor (e.g. a single SNP or a GRS) and all of its interaction partners, which do not need to be measured in the study. We treat the environment as a random effect and integrate it out, hence we require only data on the outcome and the genetic factor. Even without observing E, GxE will result in differences in trait variance across the different GRS groups. The rate of change of the outcome variance allows us to infer the underlying GxE contribution to the outcome. We first explored the performance of our method through extensive set of simulations. Most violations of our model assumptions do not lead to bias or incorrect coverage of the 95% confidence interval (which would yield badly calibrated P-values).

### 3.1 Effect of non-normality of the environmental variable or the noise

First, we explored the effect of skewness and kurtosis of the environmental variable (*E*) on the estimation bias. For this set of analyses, we fixed other parameters 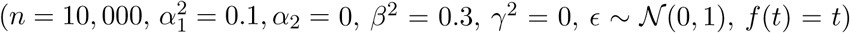 and varied only these two (*E*[***e***^3^] and *E*[***e***^4^]) in the simulations. We also explored the effect of skewness and kurtosis of the residual noise (*ϵ*) on the estimation bias in the same way but setting ***e*** ∼ 𝒩 (0, 1) and varying *E*[*ϵ*^3^] and *E*[*ϵ*^4^]. We ran simulations for all 21 possible (skewness^2^ + 1 < kurtosis) pairs of skewness (0, 1, 2, 3, 4, 5) and kurtosis (2, 3, 6, 11, 18, 27) of the environmental or noise variables and for each setting we repeated the simulation 100 times. Our results confirm that already at relatively low sample size (*n* = 10, 000) the central limit theorem ensures unbiased results for all explored non-normal distributions for both the environmental variable and the noise (Fig. 1). Similar results are obtained in case of true interaction effect (see Fig. S2 for *γ*^2^ = 0.025).

**Figure 1:**
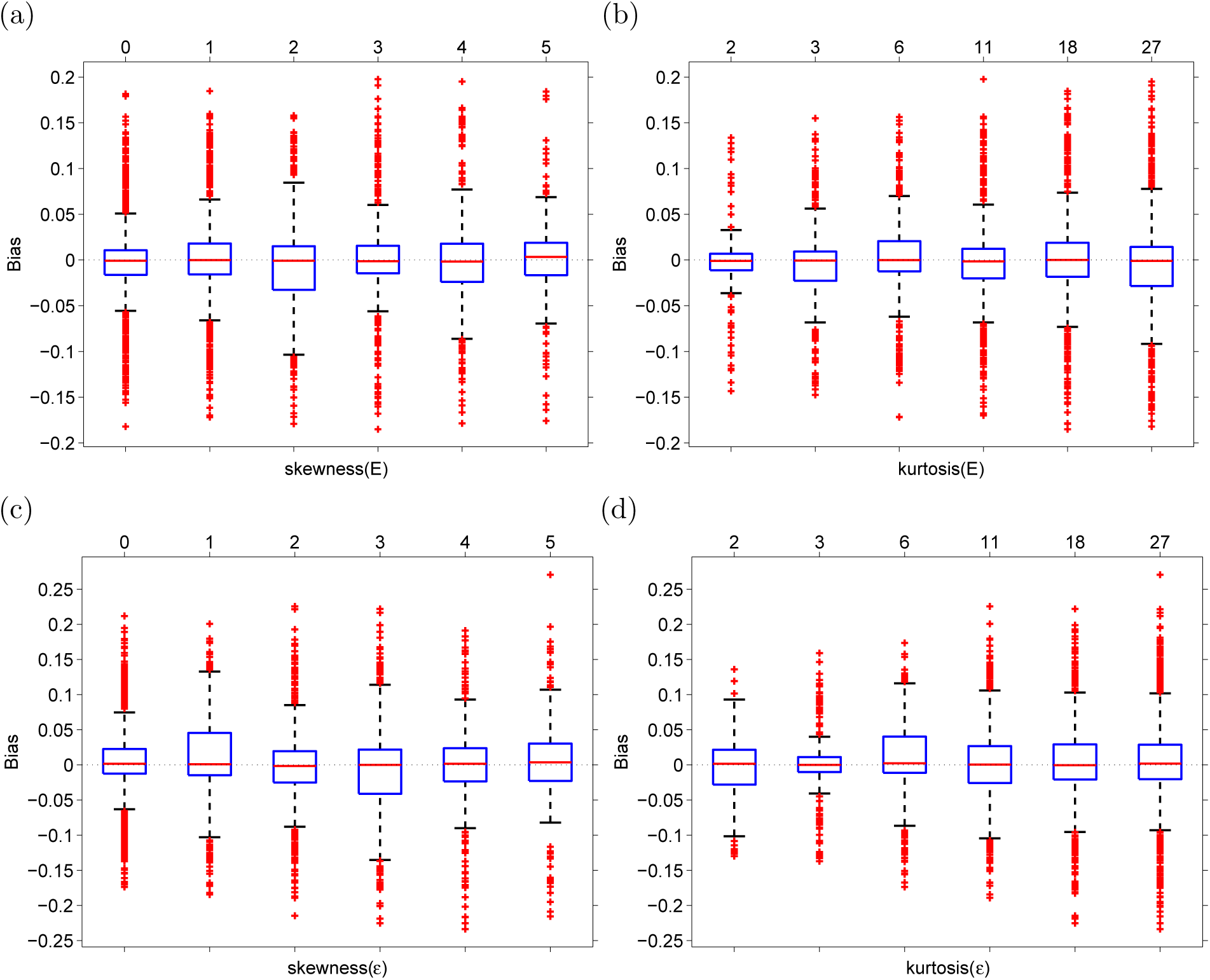
Interaction effect estimation bias as a function of skewness and kurtosis of the environmental variable (*E* - panels (a-b)) or the noise (*ϵ* - panels (c-d)). For a fixed skewness, we pooled the estimates for all possible (tested) kurtosis values and vice versa. Parameters were fixed as 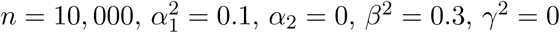.

We also observed that the bootstrap procedure (Section 2.2) ensures good coverage of the 95% confidence interval, for a wide range of distributions of the environment and noise (Fig. S3). In almost all scenarios, ∼95% of the time the true parameter fell into the 95% confidence interval. Note that for lower sample size (e.g. n=1,000) the coverage probability may deviate slightly from the nominal 95% for high kurtosis (Figure S4). The results are qualitatively identical for simulation settings when *γ* = 0 was replaced with *γ*^2^ = 0.025 (see Figure S5). Note that the root mean square error (RMSE) and power change with skewness and kurtosis of both the environment (Fig. S6) and noise (Fig. S7).

### 3.2 Outcome modelled on transformed scale

Modelling the outcome variable on a transformed scale can introduce bias in the estimation of the interaction effect size on the original scale. Since the bias is dependent on the transformation function and on the true interaction effect size, it cannot be reliably estimated in most situations. It is critical, however, to be able to distinguish between null and non-null interaction effects. For this we generated data with *γ* = 0 and *γ*^2^ = 0.05, while fixing other parameters as 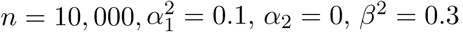. We then transformed the outcome (*Y*) using power transformations (*f* (*t*) = *t*^*p*^) with different exponents (*p* = 0, 1, 2). As described in the Methods (2.3) we also generated a fake GRS (fGRS) as a control to see how specific the observed GRS interaction is and to what extent it is due to scale/transformation effect. Results show (Fig. 2a,e) that even when the true interaction effect is null, but the outcome is transformed there is an apparent (counterfeit) interaction effect, which is negative for concave and positive for convex transformations. The fGRS analysis can, however, help distinguishing between true and counterfeit interactions by revealing whether the interaction is specific to the GRS itself. If the fGRSxE interaction estimates are close to those from the GRSxE analysis, the interaction is likely due to non-linear effects of the GRS on the scale of the observed outcome. In case of true non-zero interaction (Fig. 2b,d,f), the interaction effect estimates are also biased (except when *p* = 1), but they are significantly different from those obtained from the fGRS analysis. These results indicate that— in the case of transformed outcome—comparing interaction effect estimates coming from data with true *vs* counterfeit GRS can clearly distinguish null *vs* non-null interaction effects in most tested outcome transformation scenarios. Still, extreme transformations (*f* (*t*) = *t*^3^, see Figure S8) or large kurtosis (> 11) or skew (> 3) lead to discrepancy between 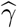 and 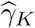 under the null. Note however, that such large kurtosis and skew values are very rare for real data (see Table S1 for 32 traits in the UK Biobank). To investigate whether inverse normal quantile transformation (INQT) of the observed outcome can improve the estimations we applied the method to INQT versions of the outcome (Fig. S9). These results show that when the skewness and kurtosis of *E* (or *ϵ*) is similar to the skewness/kurtosis of a Gaussian random variable upon applying the scale transformation function (*f* (*t*)), INQT of the outcome before applying our method can reduce type I error rate and increases power. For example when *E* (or *ϵ*) has positive skewness and *f* (*t*) is convex (e.g. *t*^2^), INQT is beneficial, because the transformation that renders the outcome normally distributed reasonably agrees with the inverse of the scale transformation (*f* ^−1^). On the other hand when positive skewness is combined with a concave transformation (e.g. log(*t*)), INQT increases type I error and decreases power at the same time.

**Figure 2:**
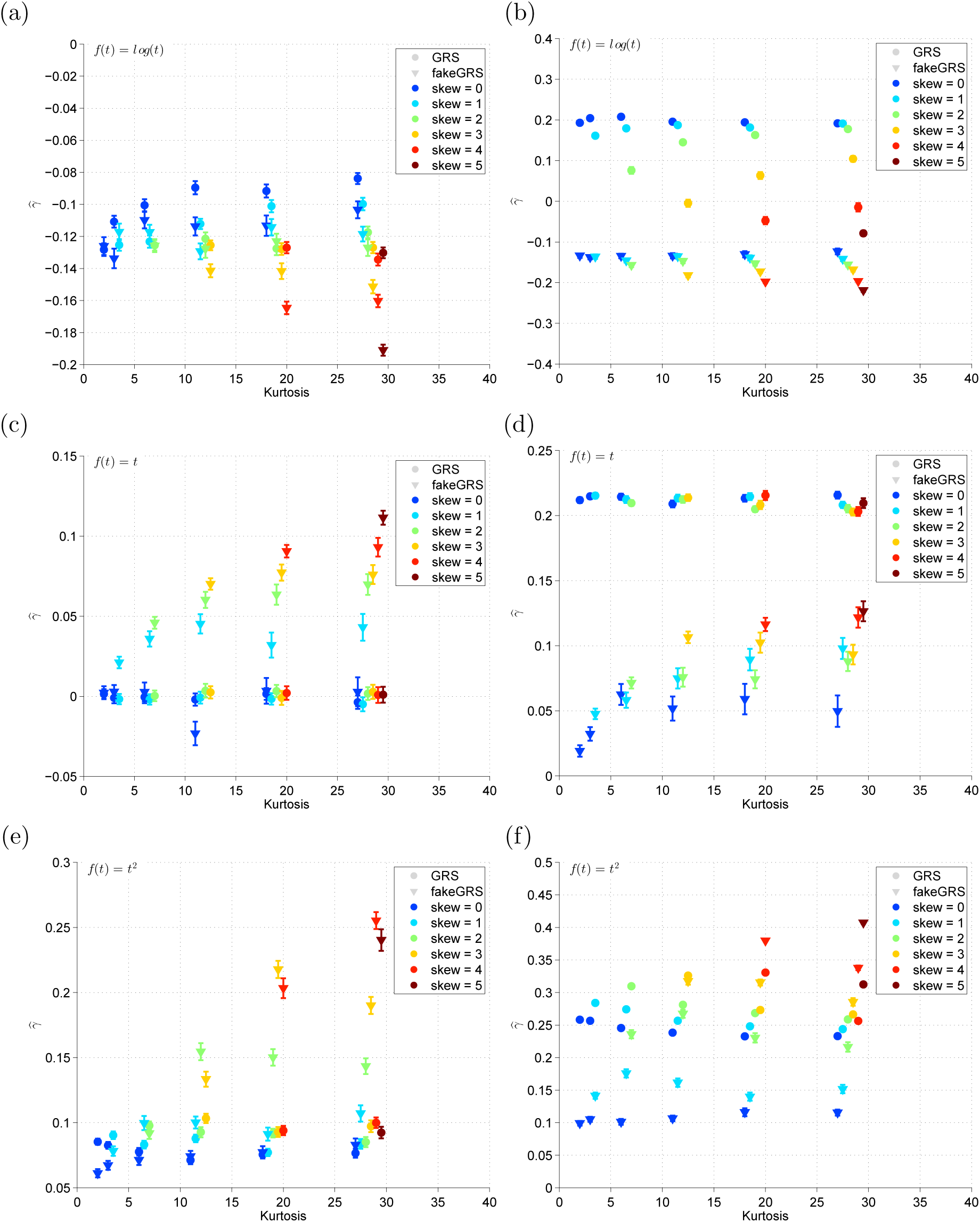
Error bars (mean± SE) for the interaction effect estimates 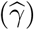 for the real data (•) and for fake *G* (∇) as a function of transformation power (*p* ∈ {0, 1, 2}: *f* (*t*) = *t*^*p*^) (rows 1-3). Other parameters were fixed at 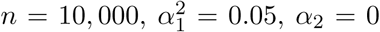, and *β*^*2*^ = 0.3. **Left column** without GxE interaction (*γ* = 0), **Right column**: With GxE interaction 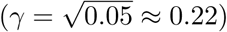.

Note that even if there is no transformation (*p* = 1) but a true interaction effect exists for the GRS, the interaction estimate for the GRS imitation will be non-zero (Fig. 2b) because the fGRS is – by construction – somewhat correlated to both *G* and *E*, hence it is bound to show some interaction. We cannot control for it as *E* is unobserved, hence cannot be regressed out. Finally, we also performed simulations mimicking UK Biobank BMI and leg impedance data and found good discriminatory power between null and true interaction scenario (Fig. S10).

### 3.3 Power analysis

Application to real data (Section 3.5) suggests that GRSxE interactions contribute ∼ 0.1%–2% of the GRS. Therefore we explored the range of *γ*^2^ ∈ [0.002, 0.02] and realistic GWAS sample sizes (*n* = 10, 000–100, 000). The simulation experiment showed that studies with sample size of *n* = 20, 000 are well-powered (> 80% power at *P* < 10^−3^) to detect interaction effects (*γ*^2^) of at least 2% and applying it to (currently considered as) large studies *n* = 100, 000 we can detect effects as low as 0.005. We have also compared the power of our method to simple linear regression interaction models when *E* is observed. The test statistic for the latter is between 4.5–6 times larger than for our method with unobserved *E* depending on skewness and kurtosis of *E* (Fig. S11). This means that as long as we observe a surrogate environment that has > 0.22 correlation with the true *E* we have more power to detect the interaction with the GRS via a linear interaction model including the proxy for *E* than using our method without any *E* observed.

### 3.4 Impact of imprecise GRS estimation

Our results showed (Fig. S12) that for both GRS estimations, at relatively stringent P-value thresholds (*P* < 10^−3^) using the estimated GRSs yields unbiased interaction effect estimates and inclusive GRSs (including SNPs also with mild P-values) lead to conservative estimates. For this reason in all other simulation in this paper we assume the GRS to be known, since it gives indistinguishable estimates for *γ* from those where the GRS is estimated from the data. Also, these calculations confirm that although our GRSs for traits in the UK biobank were estimated from the same data, given the stringent P-value threshold (5 × 10^−8^), the GxE estimates are not affected by the fact that the GRS coefficients are slightly overestimated.

To re-confirm these observations using real data, we split the filtered UK Biobank sample into two subsets (*n*_1_ = 188, 827, *n*_2_ = 188, 826). We used the first set to select 110 SNPs with (strictly) genome-wide significant level (*P* < 10^−8^) and estimated a GRS using within-sample effect size estimates (*GRS*_1_) and a second GRS (*GRS*_2_) using effect size estimates from the second subset. We then applied our method to both GRSs and obtained that the interaction estimates were not significantly different 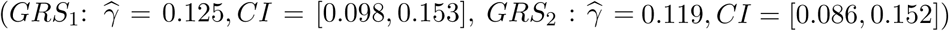.

### 3.5 Application to complex traits from the UK Biobank

We next applied our method to estimate the contribution of GxE to the heritability of complex traits. Since the interaction effects are generally modest and our method exploits variance differences, very large sample sizes from a homogeneous population are needed. For this reason we applied our method to 32 continuous complex traits measured in the UK Biobank^22^. Previous GxE studies indicated that the contribution of GxE compared to marginal genetic effects is very modest (less than a quarter of the explained variance of the marginal model). Due to this fact, we only used traits for which the GRS (built from genome-wide significant independent SNPs) explained at least 2% of the respective trait variance. We declared a trait to have significant GRSxE contribution if the interaction effect estimate could be rejected to be zero with a P-value < 1.5 × 10^−3^(= 0.05*/*32). To be conservative, when > 10% of the bootstrap estimates ended up on the boundary (i.e. Var(*ϵ*) = 0), the trait was not analysed further. Such situations may emerge when there is no interaction effect, hence *β* cannot be estimated or the likelihood surface was difficult to navigate. The estimates (summarised in Table 1 and visualised in Fig 4) revealed several interesting properties, which we describe below. In addition, we also applied the method to the inverse normal quantile transformed version of the 32 traits (see Table S2) and noticed that apart from one exception (waist-to-hip ratio) the fake GRS gave close to zero interaction effect with large variance. In addition, traits had convergence issues (maximum at the boundary) on average 88% of the starting points, which indicates that so strong transformation leads to data that is not compatible with the underlying interaction model.

**Table 1:** Estimated contribution of GRSxE effects are shown for 22 (of the 32) continuous traits measured in the UK Biobank for which no maximum likelihood estimation convergence issues were detected. Column label abbreviations are as follows: *α*_1_: *GRS* effect, *α*_2_: *GRS*^2^ effect, *β*: environmental effect, *γ*: interaction effect, *γ*_*K*_: interaction effect of fake *GRS, P*_Δ_: P-value for testing *γ* = *γ*_*K*_. Note that significantly non-zero interactions are claimed only when the P-value of the estimate 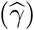 is below 0.05/32, i.e. *P*_*γ*_ < 1.5 × 10^−3^.

| Trait | $\hat{\alpha}_1$ | $\hat{\alpha}_2$ | $\hat{\beta}$ | $\hat{\gamma}(P)$ | $\hat{\gamma}_K(P)$ | $P_\Delta$ |
| --- | --- | --- | --- | --- | --- | --- |
| Body mass index (BMI) | 0.232 | 0.013 | 0.65 | 0.139 (4.3e-78) | 0.126 (5.2e-44) | 2.8e-01 |
| Trunk fat-free mass | 0.310 | 0.007 | 0.32 | 0.120 (9.0e-53) | 0.193 (<1e-300) | 1.3e-15 |
| Whole body fat-free mass | 0.301 | 0.007 | 0.39 | 0.119 (5.4e-43) | 0.191 (8.5e-37) | 3.7e-05 |
| Whole body fat mass | 0.225 | 0.011 | 0.72 | 0.124 (3.5e-42) | 0.117 (6.9e-26) | 6.3e-01 |
| Leg fat mass | 0.217 | 0.015 | 0.68 | 0.161 (3.6e-36) | 0.149 (3.0e-36) | 5.0e-01 |
| Whole body water mass | 0.301 | 0.007 | 0.39 | 0.125 (3.8e-31) | 0.190 (2.4e-10) | 4.2e-02 |
| Waist circumference | 0.198 | 0.011 | 0.65 | 0.099 (1.7e-30) | 0.111 (4.6e-24) | 3.9e-01 |
| Hip circumference | 0.221 | 0.010 | 0.69 | 0.119 (8.3e-29) | 0.146 (2.6e-75) | 4.8e-02 |
| Arm predicted mass | 0.270 | 0.006 | 0.41 | 0.117 (9.3e-29) | 0.203 (<1e-300) | 3.0e-13 |
| Weight | 0.257 | 0.011 | 0.69 | 0.123 (2.2e-26) | 0.122 (1.8e-45) | 9.3e-01 |
| Arm fat mass | 0.217 | 0.017 | 0.70 | 0.180 (3.6e-25) | 0.140 (8.4e-26) | 6.3e-02 |
| Arm fat percentage | 0.204 | 0.006 | 0.26 | 0.108 (1.1e-24) | 0.148 (1.5e-109) | 1.5e-03 |
| Trunk predicted mass | 0.309 | 0.007 | 0.32 | 0.120 (2.2e-24) | 0.195 (<1e-300) | 4.1e-09 |
| Leg fat-free mass | 0.276 | 0.008 | 0.54 | 0.126 (2.6e-23) | 0.179 (2.6e-180) | 1.9e-04 |
| Arm fat-free mass | 0.271 | 0.006 | 0.41 | 0.120 (3.1e-23) | 0.199 (2.8e-13) | 8.1e-03 |
| Basal metabolic rate | 0.287 | 0.009 | 0.48 | 0.136 (4.8e-23) | 0.187 (1.5e-22) | 2.9e-02 |
| Leg predicted mass | 0.275 | 0.008 | 0.56 | 0.124 (4.2e-11) | 0.181 (3.3e-24) | 2.7e-02 |
| Impedance of leg | 0.255 | -0.001 | 0.08 | 0.070 (1.4e-07) | -0.162 (9.2e-176) | 1.7e-57 |
| Leg fat percentage | 0.203 | 0.005 | 0.45 | 0.064 (7.7e-07) | 0.108 (2.5e-36) | 5.1e-03 |
| FVC | 0.235 | 0.008 | 0.56 | 0.071 (3.1e-05) | 0.189 (1.9e-141) | 2.2e-10 |
| Sitting height | 0.357 | 0.001 | 0.09 | 0.059 (2.9e-03) | -0.258 (<1e-300) | 1.7e-55 |
| Impedance of whole body | 0.264 | 0.003 | 0.29 | 0.048 (8.2e-03) | 0.141 (1.0e-131) | 8.1e-07 |

**Figure 3:**
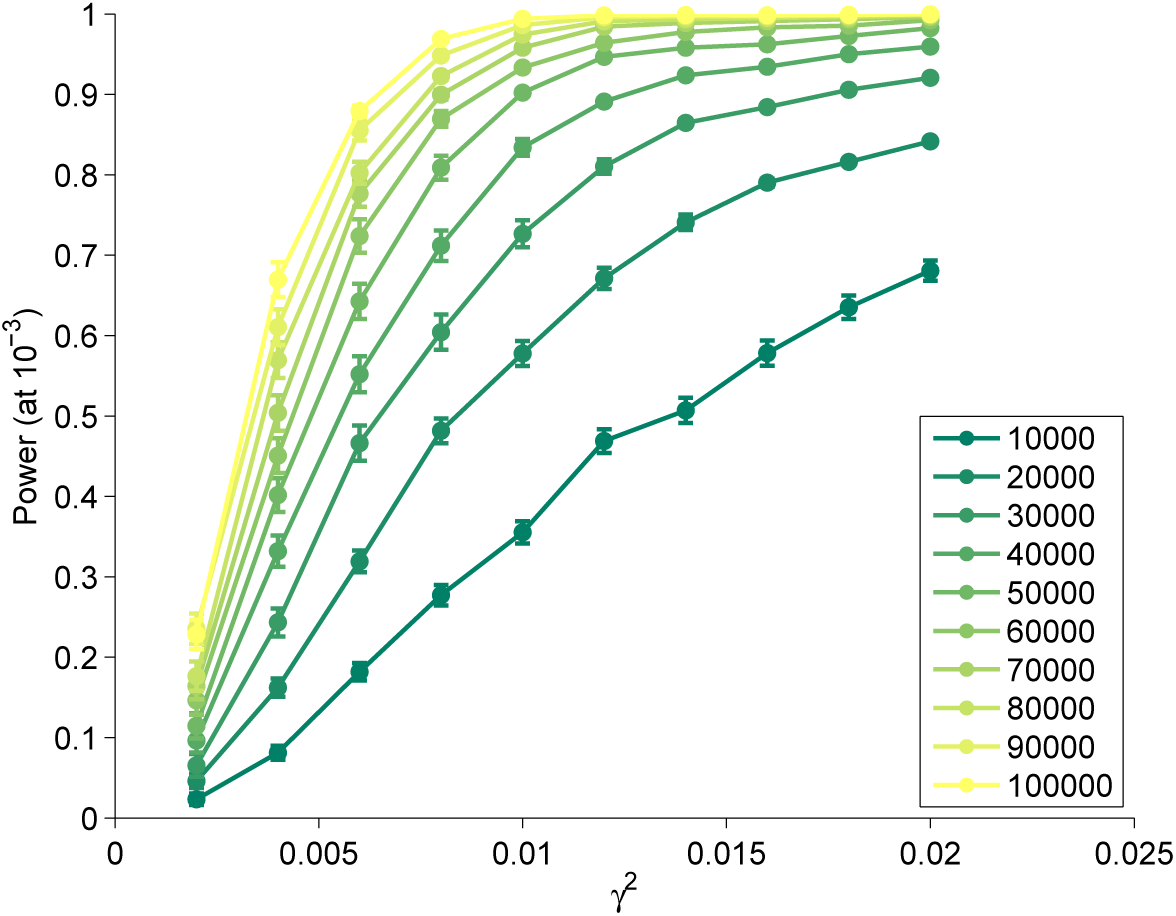
Power curves (at P-value threshold of 10^−3^) are shown for various sample sizes (10k-100k) for a range of plausible interaction effects (*γ*^2^ ∈ [0.2%, 2%]).

**Figure 4:**
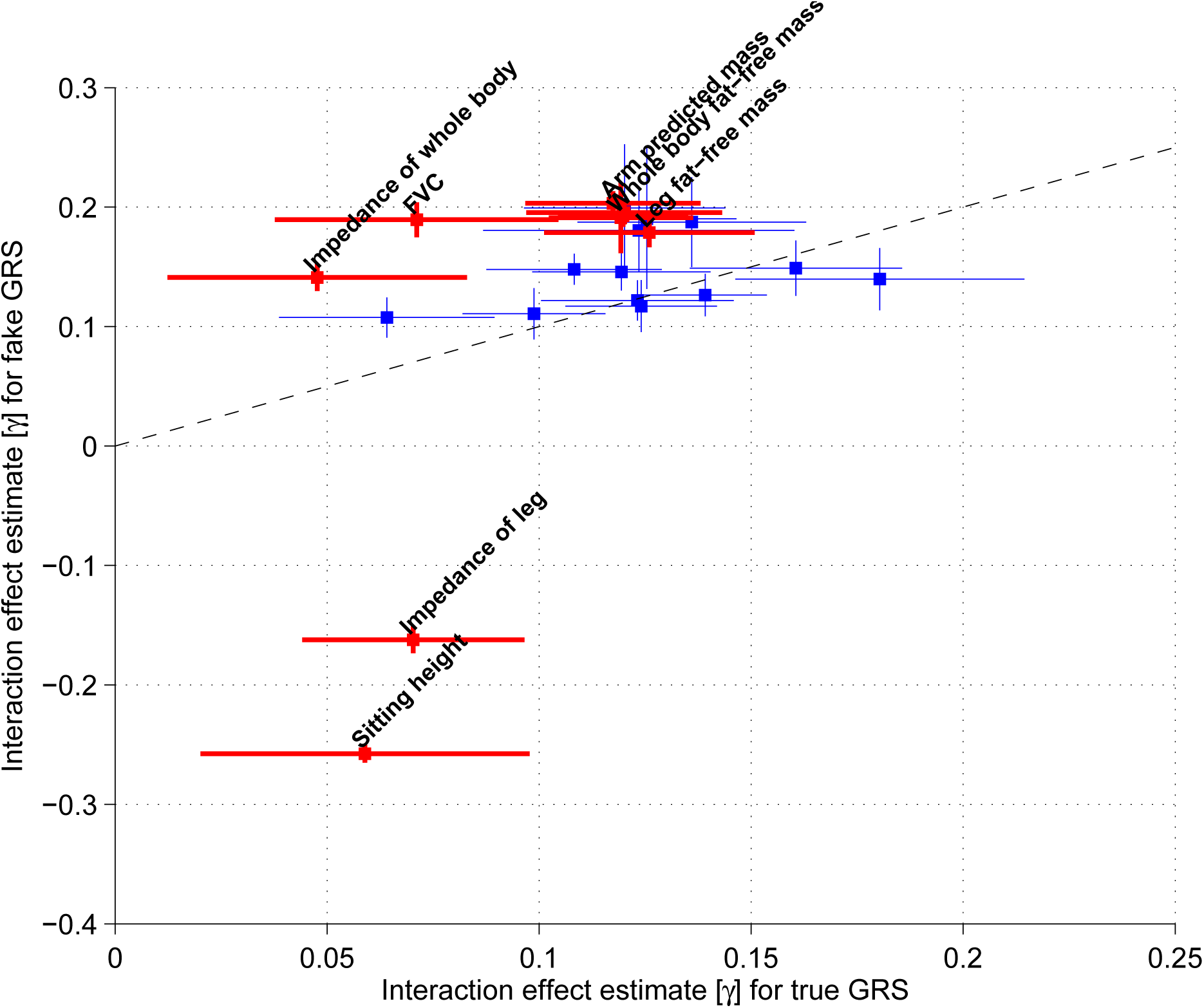
Estimated GRSxE interaction effects are shown for 22 continuous traits measured in the UK Biobank. The x-axis position of each dot (trait) represents the GxE estimate for the GRS, while the y-axis location corresponds to the interaction effect for the counterfeit GRS. Bars indicate 95% confidence intervals, those in red remain significant after Bonferroni correction at 5% type I error level when testing the difference between the two estimates. Labels for trunk predicted mass, trunk fat-free mass were omitted as they overlap with arm predicted mass and whole body fat-free mass, respectively.

While BMI shows the strongest interaction effect 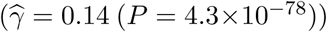, the interaction is not specific to the GRS and any similarly correlated variable would exhibit comparable interaction effect 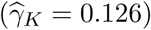. The similarity is visible when comparing the relationship between the GRS (or fGRS) and the residual variance of the outcome in the respective GRS subgroup (Fig. 5a). This means that the heteroscedastic nature of BMI is most likely due to a transformation and not driven by GxE. Interestingly, we also obtain that the unobserved “interaction partner”(E) explained 42% 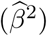 of BMI variance and the quadratic GRS significantly (*P*_*α*_2__ = 4 × 10^−40^) contributes beyond the linear term, reflecting either correlated *G - E* or true quadratic *G*^2^ *– Y* effect or transformed trait. It is important that the generated counterfeit GRS mimics not only the *GRS*-*Y* correlation, but also the non-zero *GRS*^2^-*Y* correlation. A fake GRS not fulfilling both properties resulted in almost doubled counterfeit interaction effect 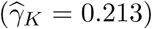. To lend further credibility to our finding, we explored the total GxE contribution once BMI is adjusted for previously reported G - alcohol intake frequency interaction in the UK Biobank^13^. Our method (re-)estimated the global uncorrected GxE contribution 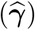, decreasing it by 0.15%, as expected. Similarly, Townsend deprivation index (TDI) - GRS interaction explained 0.09% of BMI^14^ and once BMI is adjusted for this specific interaction, our method yielded an interaction estimate reduced by 0.11%.

**Figure 5:**
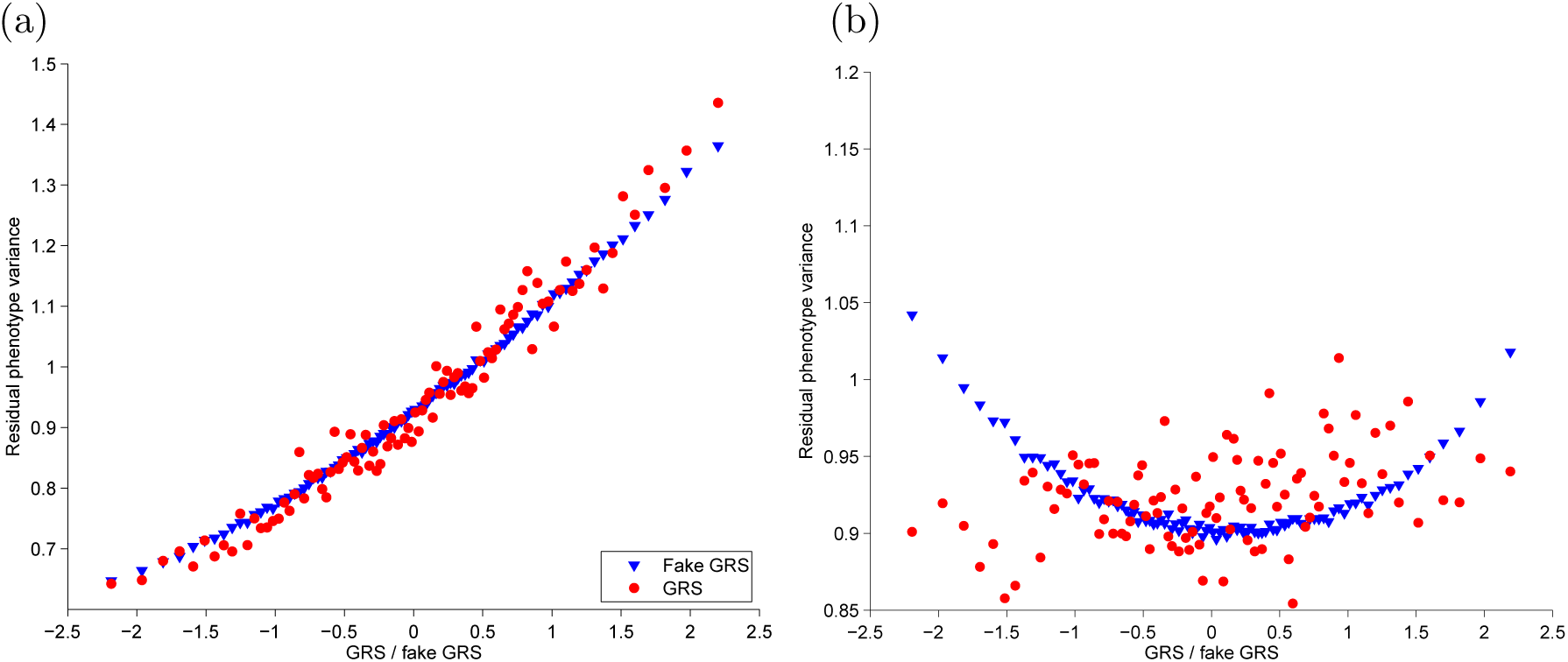
UK Biobank participants were binned into 100 equal sized groups according to their GRS values. In each group the variance of the residual outcome (Var(*Y - α*_1_*G - α*_2_*G*^2^)) was calculated and plotted against the mean GRS value in the group. Similar procedure was followed for 50 fake GRSs and the obtained variances for each bin were averaged out over the 50 repeats. Panels shows the comparison for BMI (a) and leg impedance (b).

While for the majority (59%) of the 22 traits the interaction effect of the GRS was not specific, 9 traits yielded significant difference between the interaction estimate and that for a counterfeit GRS (see Table 1 and Fig. 4) indicating a true non-null interaction. The strongest difference was observed for leg impedance where fGRS shows negative interaction 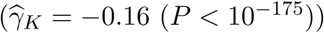 in sharp contrast to the actual GRS, which shows a positive and significant interaction 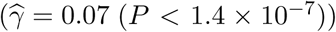. The result is a consequence of a very different mean-variance relationship for the true GRS and the fake GRS as shown in Figure 5b. Similar situation was observed for sitting height: borderline significant positive GRS interaction *vs* strong negative interaction for the counterfeit GRS. These large negative interaction effects with the counterfeit GRS are indicative that possibly the observed trait is a result of a concave transformation of an underlying variable on which the GRS and *E* act linearly.

For trunk fat-free mass the GRS shows pronounced interaction effect 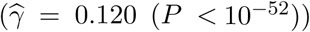, however the counterfeit GRS reveals a significantly (*P* = 1.3 × 10^−15^) stronger interaction effect 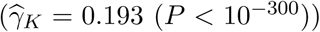. This indicates that the observed heteroscedasticity is due to the observed trait being a result of a convex transformation and on the untransformed scale the GRS would have a negative interaction. Forced vital capacity (FVC) and both arm and trunk predicted mass show a similar pattern, whereby the fGRS yields close to double sized interaction effect as the true GRS.

We further explored why the observed GRSxE interaction was not specific to the GRS in case of BMI. For this we applied our method to each constituting SNP of the GRS to obtain 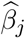 and 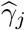 for SNP *j* of the GRS. We then established the relationship between marginal- and interaction effects driven by the general heteroscedasticity of BMI as explained in Section 2.3. It was noted that for the 376 BMI-associated SNPs the estimated 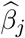 and 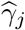 values were highly correlated (*r* = 0.27), but upon correction for overall heteroscedasticity the correlation was substantially reduced 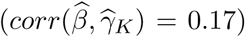 and only two SNPs survived Bonferroni correction (*P* < 0.05*/*376) (see Figure 6). Similar plot of leg impedance is shown in Figure S13.

**Figure 6:**
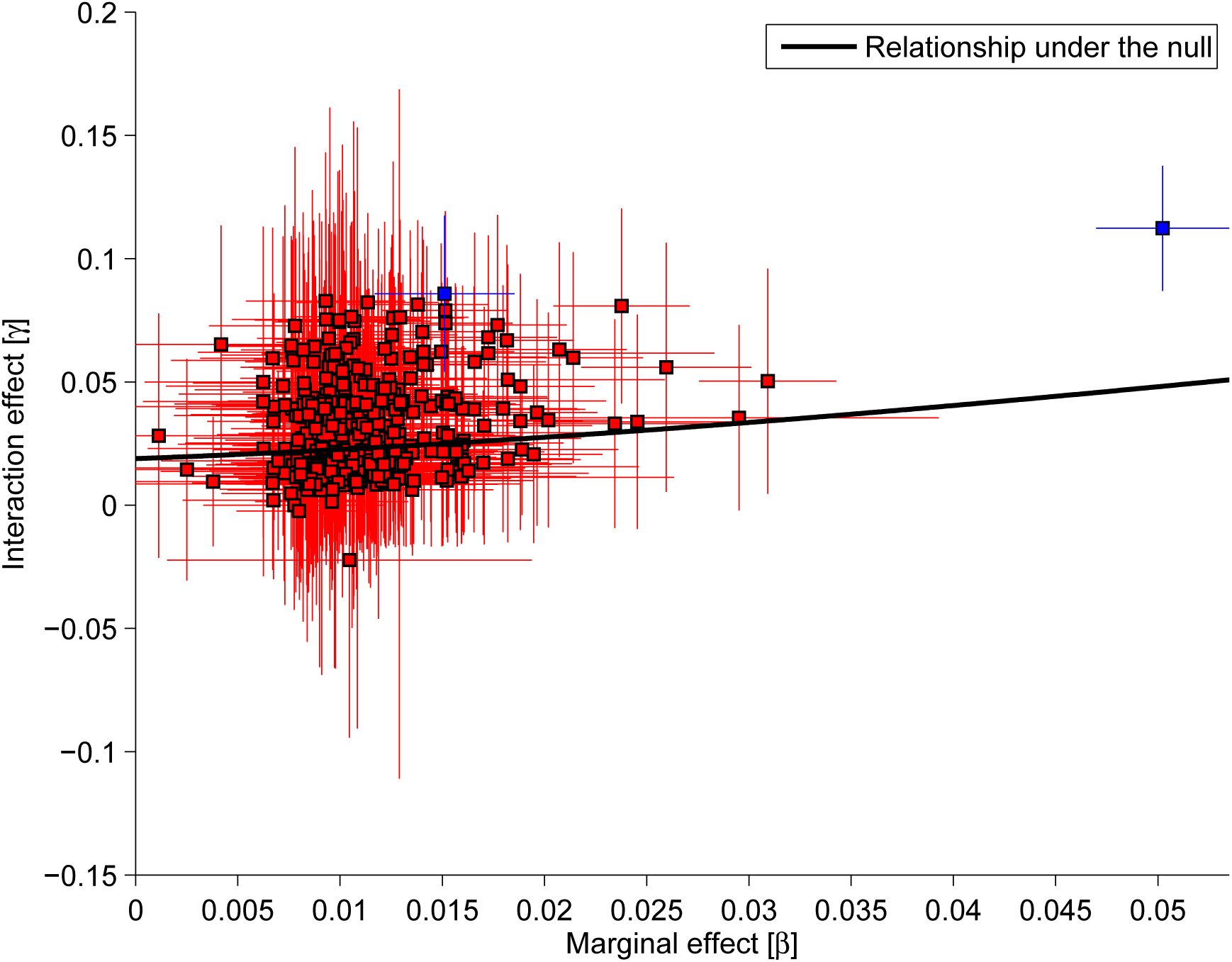
Interaction vs marginal effects for 376 BMI-associated SNPs. Black line represents the relationship for a non-specific variable, i.e. a variable with 0.05 marginal effect is expected to show an interaction effect of size ≈ 0.04. Only two of the SNPs show greater than expected interaction effect.

## 4 Discussion

We have proposed a maximum likelihood-based method to infer the extent of the total GxE interaction between a genetic risk score (which could be composed of a single SNP) and the combination of all possible continuous environmental variables. Our method is designed for a continuous outcome, but it can be extended to binary traits (see Supplementary Information 1). However, the interaction effect estimates strongly depend on the choice of the fitted link function and therefore not suitable to reliably detect interactions (data not shown). Unlike most GxE methods, it does not test a specific environment, but infers the extent of the joint contribution of all factors, while requiring none of them to be observed or measured. Since many different environmental factors potentially interact with a genetic risk score, even if some of them are binary, the (optimal) linear combination that collects all interaction partners is likely close to continuous. Thus, our modelling assumptions remain rather general. Also, testing individual interaction partners in isolation may give a false confidence of a specific modifier effect, when the tested environmental factor may simply be correlated to the true modifier. Hence, the tested and significant interaction partner may not be as specific as one might think. Similarly, some *G* × *E*_1_ may represent *G* × *G*_1_ or *E* × *E*_1_ interactions, where *G*_1_ is a genetic factor associated with *E*_1_ and *G* with environment *E*, respectively. Still, such identified GxE is an informative starting point for further investigations narrowing down the most plausible *E*.

We have shown that our approach—despite its derivation relying on the normality of both the environmental factor and residual noise—provides unbiased estimates and accurate coverage probabilities of the 95% confidence interval for a wide range of environmental and noise distributions. Additionally, we can also estimate the explained variance of the global interaction partner. Furthermore, our power analysis found that in modern biobank-sized studies (*n* > 100, 000) our method is well-powered to detect GRSxE contribution even as low as 0.5%, but this is still 2.5 times larger than the largest detected interaction for an FTO variant and alcohol consumption frequency^13^. Therefore, even in such large data sets, our method is underpowered to detect interaction between an environmental factor and individual SNPs. Its primary use is therefore for GRSxE interaction, which does implicitly assume that SNPxE interaction effect sizes are proportional to the marginal effect of the SNPs and only captures the contribution of an environmental variable interacting with many SNPs of the GRS. Others have explored evidence for trait variability at the single SNP level, but found very little evidence for it^18^. They also attempted to account for inherent mean-variance relationship of heteroscedastic traits (such as BMI), although their approach assumed that the majority of SNP should show no real interaction effect and the function of the mean-variance relationship was very simplistic, which may not be accurate for SNPs (like the one near *FTO*) with larger effects. Our approach to control for transformation-induced heteroscedasticity instead tested for the interaction effect expected for an artificial GRS with similar correlation with the outcome as the true GRS.

Recently, a similar (SNP-by-SNP) variance QTL analysis was performed for 13 traits in the UK biobank and found several associations using the Levene test^19^. However, they did not attempt to apply any correction for general heteroscedastic effects and the test is only applicable to discrete predictors, not for GRS. In the general setting variance QTLs can be detected using Double Generalised Linear Models (DGLMs)^23^. We, however, used a specific model (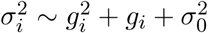 relationship) that assumes that variance QTLs are given rise entirely by a simple linear GxE model. This way, we could assess their total contribution to complex trait heritability. Assessment of the overall genomic contribution to variance modulation has been proposed^24^, however due to its computational complexity, it may not be suitable for large human population cohorts.

For main effect GWAS the outcome is often inverse normal quantile transformed (only the ranks are kept and values are replaced by corresponding standard normal quantiles). We believe that this is not necessarily the right approach for GxE interaction analysis, as such transformation will also introduce bias (usually towards the null). To demonstrate this we additionally applied our method to INQT simulated traits and observed improvements only when the INQT happened to be similar to the inverse of the scale transformation (*f* ^−1^). We have also found that when applying INQT to the 32 UK Biobank traits, in 88% of the case there were convergence issues encountered, implying that such strong transformation yields data that is incompatible with the tested model. In general, there is a key distinction in the motivations for trait transformation. One might choose to transform the trait in the hope that the residual noise will become more normally distributed and hence the P-values well-calibrated, however in case of GxE interaction models, the aim is to obtain a trait on which the genetic and environmental effects act as linearly as possible. In our view, the latter is far more important for large data sets, where the normality of the test statistic is ensured by the central limit theorem even in case of non-Gaussian residuals. In addition, our bootstrapping procedure ensures valid confidence intervals regardless of noise distribution.

Applying our method to complex traits from the UK Biobank revealed that modelling un-transformed BMI would point to substantial GRSxE contribution (increasing the 5% explained variance of the GRS by 2%), however this contribution is not specific to the GRS, any variable with comparable effect on BMI would yield very similar interaction effect. For example, log(BMI) or INQT(BMI) show no interaction effect. In general, if any two variables that are correlated with an outcome also show an interaction effect, we suspect that it is due to the trait being observed on a transformed scale and hence an observed interaction effect may not be specific. Instead of trying to guess the underlying transformation, we rather estimate how much interaction effect is attributed to a non-specific correlate of the outcome, mimicking the original risk factor. However, published GxE studies do not correct for this phenomenon. Note that our reported uncorrected total GRSxE contributions are typically much larger than any of the previously reported ones using specific environmental factors. In the UK Biobank, alcohol intake frequency - GRS interaction explained 0.19% of the BMI, representing less than the tenth of the global uncorrected GxE contribution of 1.93% estimated by our method.

As any method, ours has its own limitations. It requires access to the individual-level genetic and phenotypic data to be able to estimate the GxE contribution to a trait. Fitting a likelihood function can be time consuming, but even for UK Biobank-scale data it takes only a few seconds on four CPUs and a further 10 minutes to perform the 100 bootstraps. Our model assumes that the combination of the underlying environmental factors, summarised as *E*, is the same across SNPs and moreover it speculates that the interaction effect of each SNP is proportional to its marginal effect, which may be an oversimplification or even incorrect. We have, however, assessed the general mean-variance relationship for each trait and still found many traits with interaction effect sizes deviating from the expected, suggesting that the interaction effect is indeed somewhat proportional to the marginal effect, even after the general heteroscedasticity is accounted for. This supports an underlying interaction mechanism that impacts the overall genetic predisposition to obesity and not separately its constituents. When *E* or *E* are heavily skewed or leptokurtic and the trait is observed on a transformed scale, distinguishing null and true interaction scenarios becomes very hard and even our method may fail to do so. Real complex traits, however, only very rarely exhibit such extreme skewness and kurtosis. To be on the safe side, we recommend claiming non-zero interaction only when both the interaction effect estimate 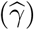 and the difference between real and counterfeit GRS (Δ) are significantly different from zero. A further limitation is that the current implementation cannot handle family data and not designed for admixed populations. If the outcome mean and variance varies across different populations, our method would interpret it as interaction, but its interpretation can be problematic. Finally, our tool is not designed to pick up interaction with non-continuous E, e.g. a G-sex interaction, which may be the most important contributor for WHR.

The proposed method could be used as a tool to establish the contribution of GxE to different traits and subsequently prioritise those with substantial global interaction effect for follow-up GxE analysis with specific environmental factors.

## Acknowledgements

This research has been conducted using the UK Biobank resource (#16389). The computations have been carried out on the HPC server of the Lausanne University Hospital. Z.K. was funded by the Swiss National Science Foundation (31003A-143914).

